# Small non-coding RNA profiling reveals inflammatory and mitochondrial related changes in aging β-cells and islet macrophages

**DOI:** 10.64898/2025.12.17.694889

**Authors:** Claudiane Guay, Jérôme Perrard, Eleonora Mangano, Véronique Menoud, Elena Aiello, Daniela Fignani, Francesco Dotta, Susan Bonner-Weir, Guido Sebastiani, Cristina Aguayo-Mazzucato, Cristina Cosentino, Romano Regazzi

**Author notes:** Corresponding authors: Prof. Romano Regazzi and Dr Cristina Cosentino Department of Fundamental Neurosciences, Rue du Bugnon 9, 1005 Lausanne, Switzerland, and. These authors contributed equally to this work (Cristina Cosentino, Romano Regazzi).

## Abstract

Aging is accompanied by functional decline and increased senescence of pancreatic β-cells. These changes may be influenced by islet-resident macrophages (iMACs) that remodel tissue in response to environmental cues. To explore the molecular mechanisms underlying β-cell aging and senescence, we profiled small non-coding RNAs (sncRNAs) in FACS-sorted β-cells and iMACs from 3-, 12-, and 22-month-old mouse islets or from senescence-associated βgal (SA-βgal) positive β-cells of 8-month-old mice. Overall, senescent β-cells displayed distinct sncRNA signatures that only partially overlapped with those of aging. However, several miRNAs previously found to be deregulated in obese or diabetic conditions were modulated in both aging and senescent β-cells, including upregulation of miRNAs linked to inflammation. In vitro exposure to pro-inflammatory cytokines partially reproduced these profiles. Aging also reshaped the β-cell tRNA-derived fragment (tRF) pool, enhancing global mitochondrial tRF levels. Interestingly, some changes in miRNAs and tRFs were β-cell specific, whereas others occurred also in other aged metabolic tissues. iMACs also showed age-related sncRNA remodeling, including upregulation of anti-inflammatory miRNAs and mitochondrial tRFs, suggesting adaptive immune reprogramming. Together, these data reveal a profound, coordinated reshaping of the sncRNA landscape in β-cells and iMACs during aging, offering new insights into molecular mechanisms driving age-related islet dysfunction.

## INTRODUCTION

Biological aging results from the accumulation of cellular damage that leads to a progressive decline in physiological functions, favoring disease development and mortality. While being a fundamental process, aging is influenced by genetic predisposition and environmental factors. With the global increase in life expectancy, understanding the complex cellular mechanisms linked to aging is crucial to favor healthy aging and reduce disease outcomes in elderly people. Aging represents a risk factor for metabolic diseases like diabetes because of the establishment of insulin resistance in insulin target tissues and the progressive decline of pancreatic β-cell ability to compensate [1–3]. Several hallmarks have been associated with aging [4] including cellular senescence, altered intercellular communication and epigenetic alterations.

Accumulation of senescent β-cells has been observed in both elderly humans and mice, and their proportion increase in response to glucose intolerance and to cellular stress [5–7]. Importantly, clearance of senescent β-cells using senolytic drugs and/or by genetic ablation prevented both type 1 (T1D) and type 2 (T2D) diabetes development in different mouse models [6, 8].

An important alteration in islet intercellular communication is the establishment of low-grade chronic inflammation [4], an aging hallmark shared with obesity, another common risk factor for T2D. Among the mechanisms possibly involved, islet-resident macrophages (iMACs) emerge as new players [9, 10]. Under physiological conditions, iMACs exhibit a unique pro-inflammatory profile that favors β-cell function, while insulin resistant conditions induce iMAC expansion and transcriptional reprogramming that exacerbates β-cell dysfunction. Increased macrophage numbers have been observed in aged mouse islets [11], however their activation phenotype remains to be investigated.

As part of age-related epigenetic alterations, recent evidence shows that the correlation between mRNA and protein expression is progressively lost during aging [12], pointing to a deregulation of the post-transcriptional regulatory network. Non-coding RNAs orchestrate a variety of post-transcriptional gene regulatory mechanisms. NcRNAs are classified based on nucleotide length: long non-coding RNAs (>200nt) and small ncRNAs (sncRNAs, <200nt). Beside well described sncRNA classes like microRNAs (miRNAs) and PIWI-interacting RNA (piRNAs), other categories of sncRNAs with specific mode of actions in gene regulation are continuously discovered. Fragments derived from the cleavage of transfer RNAs (tRFs) have been identified as functional sncRNAs binding to different protein and RNA interactors [13]. Next-generation sequencing studies revealed the existance of isoforms of miRNAs (isomiRs) that differ by one or more nucleotides from the canonical miRNA sequence, resulting in differences in target recognition or stability [14, 15]. Both tRFs and isomiRs have been identified in pancreatic β-cells and have been proposed to be involved in diabetes development [16–19]. miRNA deregulation in aging has been scarcely studied. We previously performed a microarray on aged rat islets and observed that an increase of miR-34a-5p and a decrease of miR-181a-5p were associated with impaired β-cell proliferation in elderly rat islets [20]. The contribution of other classes of sncRNAs to age-related alterations in islet function remains to be established.

Here we took advantage of small RNA sequencing technologies and refined bioinformatics pipelines to obtain a comprehensive sncRNA profiling of aging β-cells and iMACs. Aged β-cells showed changes in the level of several miRNAs that are involved in the control of β-cell functions and are deregulated in T1D and/or T2D. Globally, aged β-cells exhibited an inflammatory miRNA signature while iMACs displayed a miRNA profile reflecting the acquisition of a tissue homeostatic reparative phenotype. Both β-cells and iMACs showed increased mitochondrial tRF levels in aging, suggesting metabolic adaptation. Senescent β-cells in adult mice displayed a sncRNA expression profile partially overlapping with that of aged β-cells, with activation of inflammatory miRNA-dependent mechanisms and anti-proliferative signals. In addition to the analysis of the miRNA and tRF patterns associated with aging, we also provide data on piRNAs, isomiRs and other less characterized sncRNAs modulated during aging.

## METHODS

### Experimental models Mice

All animal procedures were performed in accordance with the NIH guidelines, and protocols were approved by the Swiss Research Councils and Veterinary Offices under the animal authorization number VD2495x4, or by the Animal Care and Use Committee from the Joslin Diabetes Center (Protocol number 2022-01). Wild-type C57BL/6J mice were acquired from Jackson Labs and wild-type C57BL/6NRj mice were obtained from Janvier Laboratories or from our colony. Mice were housed on a 12-h light/dark cycle under standard conditions with *ad-libitum* chow diet. Glycemia and body weight were all taken under fed state between 8 and 10AM.

### Mouse islet isolation and cell dispersion

Mouse islets were isolated by collagenase digestion of the pancreas [21] followed by Histopaque density gradient to separate them from digested exocrine tissue. At the end of the isolation, islets were hand-picked and incubated for a minimum of 2h at 37°C in RPMI 1640-GlutaMAX medium (ThermoFisher) containing 11.1mM glucose and 2.06 mM L-glutamine and supplemented with 10% fetal calf serum (Sigma), 10 mM HEPES, pH 7.4, 1 mM sodium pyruvate, 100 mg/mL streptomycin and 100 IU/mL penicillin. Dissociated islet cells were obtained by incubating the islets in Ca^2+^/Mg^2+^ free phosphate buffered saline, 3 mM EGTA and 0.002% trypsin for 2-3 min at 37°C.

### Fluorescence-Activated Cell Sorting (FACS) of islet cells

Once dispersed, mouse islet cells were FAC-sorted based on two different strategies as described previously to obtain 1) senescent versus non-senescent β cells [8] or 2) β-cells and islet-resident macrophages (iMACs) of aged mice [9]. To obtain senescent β-cells, dispersed islet cells from 7–8-month-old C57Bl6/J male mice were incubated at 37 °C for 1h with a fluorescent substrate of the senescence-associated acidic β-galactosidase (SA-β-Gal) (Enzo Life Sciences enz-kit 130-0010). Islet cells were then resuspended in FACS buffer containing 2% FBS (Cellgro, Manassas, VA). Using a DakoCytomation MoFlo Cytometer (Dako, Ft. Collins, CO) or Aria (BD FACS Aria IIu and BD FACS Aria Special Order Research Product), cells were gated according to forward scatter to obtain an enriched fraction of β-cells and FACS-sorted based on SA-β-Gal activity. The percentage of β-cells by insulin staining ranged from 80%–90%. Propidium iodide was used to exclude dead cells. SA-β-Gal positive (pos, +) and SA-β-Gal negative (neg, -) subpopulations were collected for RNA extraction and small non-coding RNA sequencing. Complete FACS gating strategy can be seen in Figure S1 of ref [8].

To FACS sort β-cells and iMACs from different aged C57Bl6/J mice, islet cells from four males aged of 3 months or two males aged of 12 or 22 months were pooled (around 600-800 dispersed islets per sample). Fractions enriched for β-cells were obtained based on β-cell autofluorescence [22] while fractions enriched in iMACs were sorted based on the expression of CD45, Cd11b, F4/80 and Cd11c markers. As previously described [9] dissociated islet cells were washed once with FACS buffer (0.1% BSA, 2mM EDTA, 11mM glucose in PBS) and incubated for 5 minutes with TruStain FcX™ (anti-mouse CD16/32) Antibody (BioLegend) at 4°C. Cells were incubated for 30 min in the dark at 4°C with the following antibodies: 1:200 of FITC anti-CD45, 1:100 brilliant violet CD11b, 1:100 APC F4/80 and 1:100 PE CD11c (BioLegend). Cells were then washed twice with FACS buffer and sorted by FCF-Aria-II (SORP). β-cell purity was assessed as previously described [23]. On average, β-cell fractions contained 99.1 ± 0.9% insulin-positive cells and 0.6 ± 0.6% glucagon-positive cells. FlowJo^TM^ 10 software was used for quantification of iMACs from FACS-derived data.

### Cytokine treatment of dispersed islet cells

Dispersed islet cells were seeded and cultured in complete RPMI medium. In vitro treatments were performed as shown in Fig. 5a. To mimic long term aging and senescence, dispersed islet cells were incubated with a mix of pro-inflammatory cytokines (30 ng/ml IFNγ, 10 ng/ml TNFα and 1 ng/ml IL-1β) for 48h followed by media change and a wash out period of 72h. In parallel, acute treatment with pro-inflammatory cytokines was performed in the last 48h of the culture (Fig. 5a). RNA was extracted in all conditions after a total of 6 days of cell culture.

### RNA Assays

RNA was extracted from islet cells using the miRNeasy micro kit (Qiagen). For small RNA-sequencing, cDNA libraries were prepared using QIAseq miRNA Library Kit starting from 5-10 ng of RNA and single-end sequenced using Illumina technology. Mature miRNA and tRF levels were measured using the miRCURY LNA Universal microRNA PCR assays (Qiagen) starting from 50 to 200 ng of RNA. Small non-coding RNA levels were normalized to miRNAs highly expressed in β-cells (miR-7a-5p and let-7b-5p) or macrophages (miR-16-5p), and not modified under the studied conditions based on NormFind strategy [24]. Tissue small non-coding RNA levels were normalized to let-7a-5p and let-7b-5p, whose expression was stable across the different conditions. For mRNA measurement, RNA samples were treated with DNase prior to analysis. Gene expression levels were measured by qPCR (BioRad) and results were normalized to the housekeeping gene *Hprt*. Custom primer sequences are provided in Suppl. Table 1. All qPCR measurements were performed in duplicates and expression levels were analyzed using the 2^-ΔΔCt^ method. Of note, since the amount of RNA extracted from FACS-sorted islet cells was very low, tRNA modifications that may interfere with the quantification of tRFs could not be removed from RNA samples and may therefore reduce the amount of some tRFs detected in our analysis.

### RNAseq analysis

For miRNA, piRNA and isomiR profiling, FASTQ files were uploaded on the Qiagen RNA-seq analysis portal. For tRF profiling, as previously described [17], following removal of adapter sequences, identical reads longer than 16 nts were collapsed based on Unique Molecular Identifiers and aligned to the mouse genome (GRCm38.p6). Aligned reads were mapped to the mature tRNA sequences from the GtRNAdb database (Release 1.0; http://gtrnadb.ucsc.edu/, accessed March 2025) and to the 22 mouse mitochondrial tRNA sequences from https://www.ncbi.nlm.nih.gov/nuccore/J01420.1 (accessed March 2025) using bowtie (version 1; http://bowtie.cbcb.umd.edu). Bowtie parameters were set to output only perfect matches to tRNA sequences. Reads not aligning to the chosen reference sncRNA databases and unambiguously mapping to one genomic location were considered for an unbiased identification of potential novel sncRNAs.

Upon sequence annotation, differentially expressed sncRNAs were identified using edgeR (version 4.2.2) in R packages (http://bioconductor.org). For each sncRNAs, *p-*values and false discovery rates (FDRs) were obtained based on the model of negative binomial distribution. Fold changes of gene expression were also estimated within the edgeR package. The criterion for differential expressions of sncRNAs has been set as fold change (FC) >1.5 and FDR <0.1.

### Re-analysis of RNAseq data from mouse islet treated with cytokines

To investigate if isomiRs are also deregulated in response to inflammation, we re-analyzed RNAseq fastQ files from our laboratory. Mouse islets were isolated and cultured for 24h in control conditions or treated with a mix of pro-inflammatory cytokines (30 ng/ml IFNγ, 10 ng/ml TNFα and 1 ng/ml IL-1β). RNA was extracted using the miRNeasy micro kit (Qiagen) and RNA samples were sent to Arraystar Inc (Maryland, US) for sequencing. FASTQ files obtained from the small RNA-seq experiment were analyzed with the sRNAbench online pipeline. Raw reads underwent adapter removal and PCR duplicate filtering, followed by stringent quality assessment to retain only high-fidelity sequences (Q ≥ 30). Reads were mapped to the mouse genome (GRCm39) using Bowtie with specific parameters and annotated them using miRBase release 22.1. Low-quality and ambiguous sequences were filtered, followed by a relative abundance filter based on reads per million (RPM), with sequences contributing less than 1% removed. IsomiRs were classified into different classes according to the post-transcriptional modification compared with the canonical miRNA sequence [16]. An overlap analysis was performed across the different datasets to identify shared isomiR sequences in inflammatory context.

### Statistical analysis

Data are expressed as mean ± SD. Statistical significance was determined using unpaired two-tailed Student’s t test or, for multiple comparisons, with one-way ANOVA, followed by post-hoc Dunnett test (Graph Pad Prism 10). P-values less than 0.05 (p < 0.05) were considered statistically significant.

## RESULTS

### Comprehensive analysis of the changes in the small non-coding RNA profile in aging β-cells

Pancreatic islets were isolated from young adult (3-month-old), middle-aged (12-month-old) and elderly (22-month-old) wild-type C57BL/6J male mice. Fed blood glucose levels were similar between age groups while body weight was higher in 12- and 24-month-old mice (**Suppl. Figure 1.a-b**). To obtain a comprehensive data set of the sncRNAs expressed in insulin-secreting cells, we performed a small RNA sequencing on FACS sorted β-cells from these 3 age groups (**Suppl. Figure 1.c-d**). Sequencing reads were then annotated to the mouse genome and mapped to reference databases for miRNA, isomiR, piRNA, and tRF identification. In addition, reads not matching reference sncRNAs but unambiguously mapping to one genomic location were used for differential expression analysis. This systematic and unbiased analysis revealed major modulation in the sncRNA profile of β-cells occurring during aging. Using a cut-off of Fold change ≥1.5 and FDR≤0.1, we could detect age-dependent changes in all different sncRNA biotype categories in middle-aged and elderly β-cells compared to young adults (**Suppl. Figure 1.e**).

### Differential miRNA and isomiR expression in aged β-cells

We next further characterized the modulated miRNA signature because of their crucial role in pancreatic islet homeostasis and diabetes development [25–27]. Analysis of the miRNA profile of FACS-sorted β-cells led to the identification of 60 and 176 miRNAs that are differentially expressed in 12-month-old and in 22-month-old mice, respectively, compared to young 3-month-old mice (FDR p-value <0.1, FC>1.5; **Figure 1.a-b**). Among the age-modulated miRNAs, 17 were downregulated and 21 were upregulated in β-cells of both 12- and 22-month groups (**Figure 1.c**). Notably, some miRNAs such as miR-146a-5p, miR-215-3p and miR-215-5p were found to be downregulated at 12 months but upregulated in older mice (22 months). To identify pathways potentially affected by the modulated miRNAs, we performed a functional enrichment analysis. Downregulated miRNAs in 22-month-old β-cells were associated to the regulation of FoxO and mTOR signaling (**Figure 1.d**). These pathways control many essential cellular processes, such as cell cycle, survival, metabolism and proliferation, and contribute to cellular aging by modulating autophagy and oxidative stress. Other overrepresented KEGG pathways in downregulated miRNAs were insulin resistance and autophagy, while miRTar Base Gene Ontology enrichment highlighted the downregulation of miRNAs involved in cellular responses to hypoxia and to insulin resistance (**Figure 1.e**). The annotation of upregulated miRNAs did not return any significant pathway. We then analyzed whether the miRNAs modified during aging are involved in different β-cell processes and in diabetes development. Based on a meta-analysis of the existing literature, we identified groups of miRNAs associated to islet inflammation (**Figure 1.f**), β-cell proliferation (**Figure 1.g**), β-cell maturation (**Figure 1.h**), insulin secretion (**Figure 1.i**), and T1D and T2D (**Figure 1.j-m**). As shown in the heatmap plots, miRNAs promoting islet inflammation generally increase with age, while miRNAs associated with β-cell proliferation and maturation tend to change towards a highly mature β-cell phenotype with limited proliferative capacity. Moreover, at least part of the changes detected in β-cells of elderly animals overlapped with those observed during T1D and T2D pathogenesis.

**FIGURE 1.**
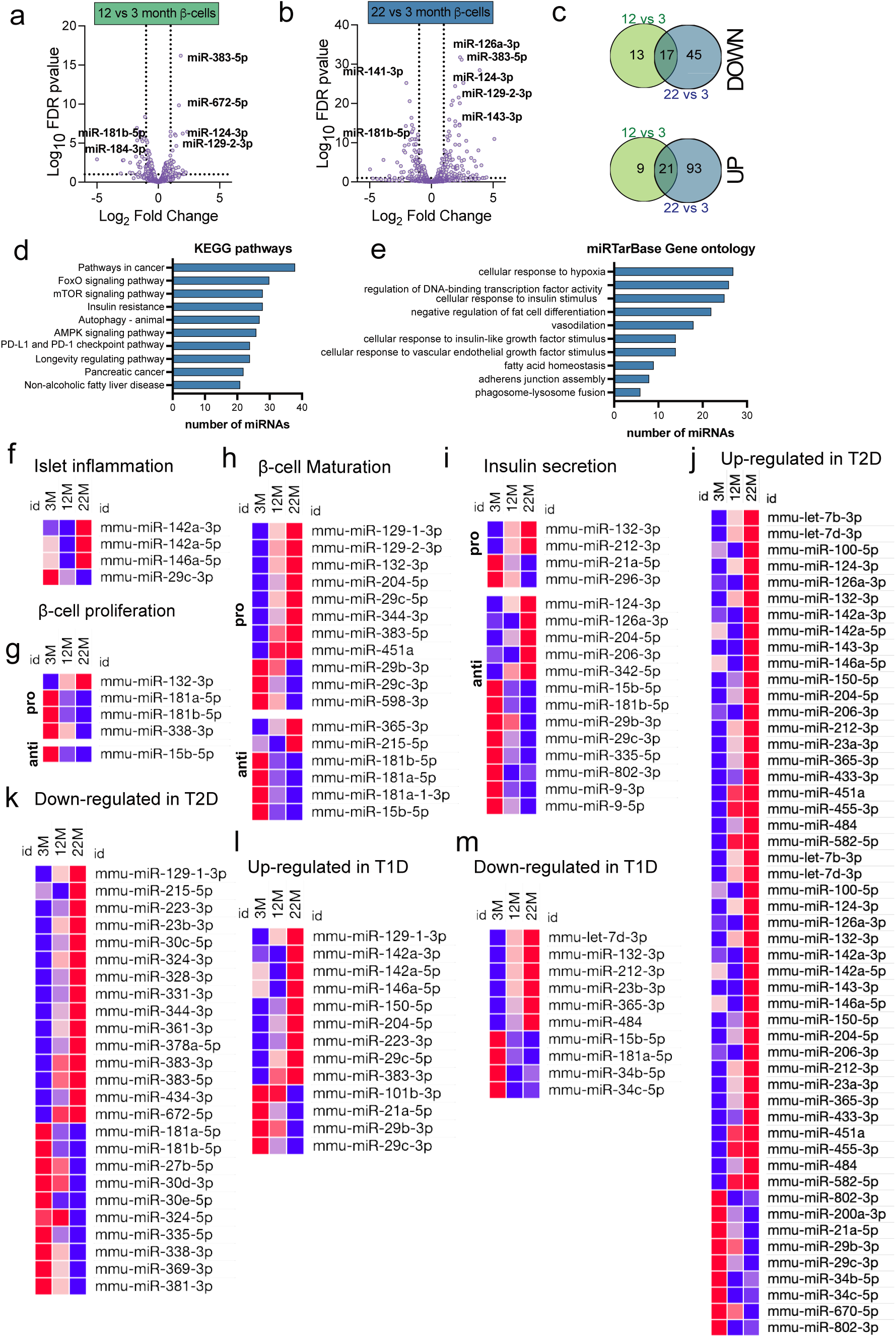
Differential miRNA expression in aged β-cells. Small RNAseq on FACS-sorted β-cells from young adult (3-month), middle-aged (12-month) and elderly (22-month) C57BL/6J male mice. a-b) Volcano Plots show differentially expressed miRNAs in 12-month (a) or 22-month (b) versus 3-month-old β-cells (FDR p-value <0.1, FC>1.5). c) Venn Diagram representation of the number of miRNAs commonly up- or down-regulated in 12- and 22-month-old β-cells. d) Overrepresented KEGG pathways and e) miRTar Base Gene Ontology enrichments associated with miRNA downregulation of 22-month-old β-cells. f-m) Heatmap plots showing miRNAs significantly deregulated in 22-month-old β-cells that were previously reported in the literature to be associated with islet inflammation (f), β-cell proliferation (g), β-cell maturation (h), or insulin secretion (i), or deregulated in islet from T2D (j-k) or T1D (l-m) models.

Beside canonical miRNAs, we also identified several isomiRs differentially expressed in aging β-cells (Figure 2.a-b). As for their canonical counterparts, the isomiRs miR-383-5p.Cs, miR-126a-5p.T, miR-126a-3p.ts.T and miR-126a-3p.A were found to be more abundant in aged β-cells. A group of isomiRs have been recently identified to be specifically expressed in β-cells compared to other islet cell types [16]. Among these, isomiRs of miR-221-3p and miR-204-5p were found to be upregulated in aging while isomiRs from miR-141-3p and miR-335-5p were downregulated (Figure 2.b). Since several miRNAs deregulated in aging are linked to inflammation, we assessed the impact of inflammatory conditions on isomiR levels. For this purpose, we performed small RNAseq from mouse islets treated with pro-inflammatory cytokines (Suppl. Figure 2.a). Cytokine treatment resulted in the down-regulation of 13 miRNAs and 46 isomiRs, and the upregulation of 10 miRNAs and 62 isomiRs (FDR p-value <0.1, FC>1.5) (Suppl. Figure 2.b). Interestingly, our sequencing data confirmed the upregulation of the inflammation-associated miR-146a-5p [28, 29] and of the T1D-associated miR-1a-3p [30] in response to cytokines. Among the isomiRs modulated by cytokines, we found that miR-143-3p.T, showing a 3’ insertion was also upregulated in aged β-cells. miR-485-3p.T which is induced in β-cell aging was also deregulated in response to cytokines, but in this case its level was downregulated (Figure 2.b).

**FIGURE 2.**
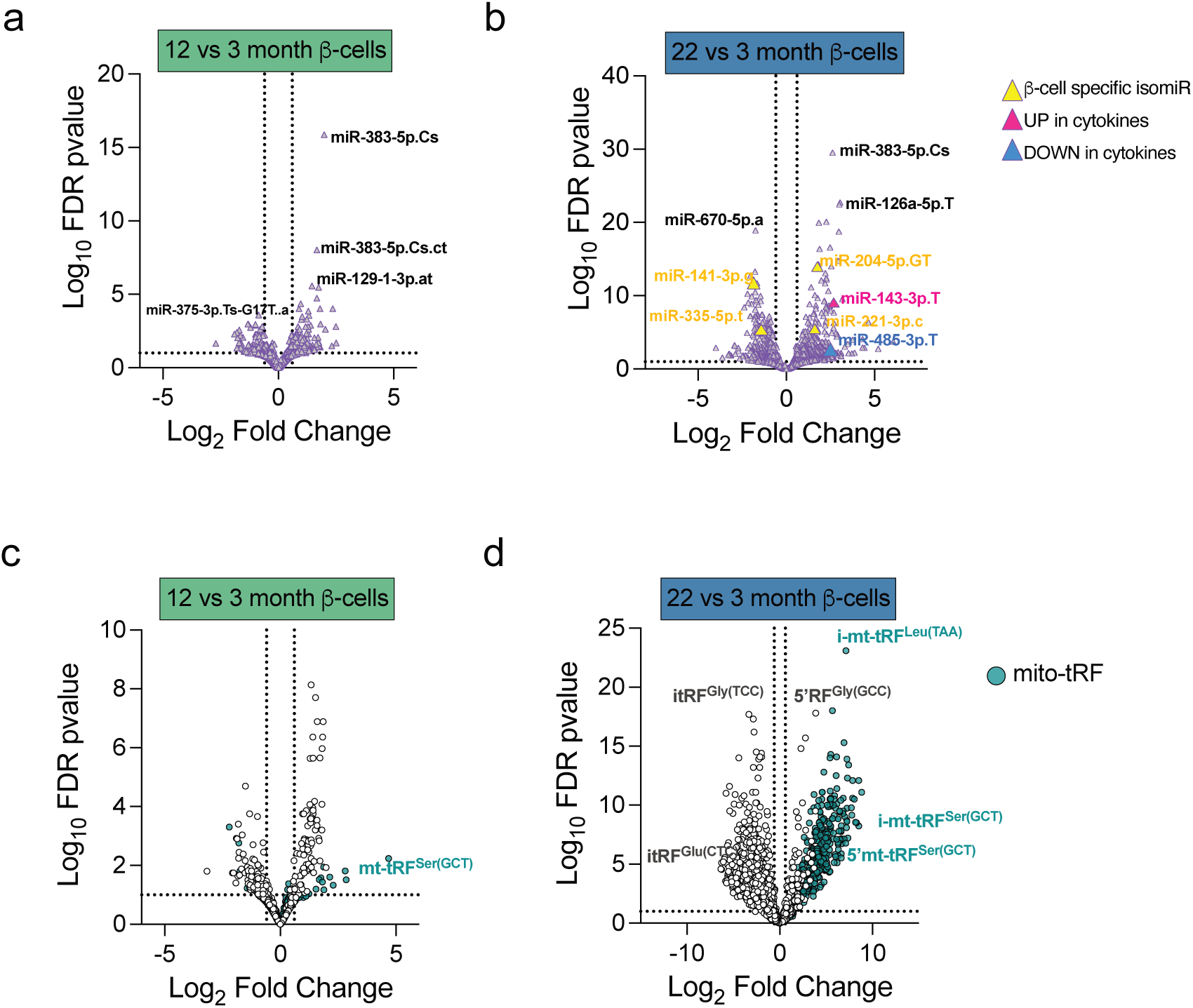
Differential isomiR and tRF levels in aged β -cells. Small RNAseq on FACS-sorted β-cells from young adult (3 months), middle-aged (12 months) and elderly (22 months) C57BL/6J male mice. a-b) Volcano Plots show differential levels of isomiRs in 12-month (a) or 22-month (b) versus 3-month-old β-cells (FDR p-value <0.1, FC>1.5). Yellow triangles highlight isomiRs previously found to be β-cell specific, and pink and blue triangles isomiRs upregulated or downregulated, respectively, in islets treated with pro-inflammatory cytokines. c-d) Volcano Plots show differential levels of tRFs in 12-month (a) or 22-month (b) versus 3-month-old β-cells (FDR p-value <0.1, FC>1.5). White circles correspond to tRFs derived from nuclear-encoded tRNAs and turquoise circles refer to tRFs derived from mitochondrially-encoded tRNAs.

### Impact of aging on the β-cell tRF profile

tRNA cleavage generates sncRNA fragments playing an important role in islet cell homeostasis. We further analyzed the nature of modulated tRFs because of their emerging role in gene regulation and cellular function in islet cells [17, 18, 31]. β-cell sequencing revealed an important age-dependent modulation of the tRF pool. Indeed, compared to 3-month-old mice, 190 tRFs displayed changes at 12 months and 1543 at 22 months (**Figure 2.c-d, Suppl. Figure 1e**). Interestingly, several mitochondrially-encoded tRFs were upregulated in 22-month-old β-cells (**Figure 2.d**). The upregulated mitochondrial tRFs included the internal tRF derived from mt-tRNA^Leu(TAA)^, which we previously showed to regulate mitochondrial respiration and glucose-induced insulin secretion and 5’tRF^Ser(GCT)^, which is repressed in islet cells from 16-week-old db/db mice [31]. To assess whether the increase in mt-tRFs reflects a global increase in the mitochondrial mass, we measured the level of the mitochondrially-encoded ribosomal RNAs 16S and 12S. These mitochondrial RNAs were not significantly affected in aging mice, although we observed a tendency to increase with age (**Suppl. Figure 3.a-b**). Moreover, the unbiased analysis highlighted an age-dependent overexpression of sequences mapping to mitochondrial RNA transcripts, such as ribosomal RNAs and mitochondrially encoded protein-coding genes (**Suppl. Figure 3.c-d)**. The settings of our small RNA sequencing allowed the detection of sequence lengths up to 80 nucleotides. This allowed us to detect sequences of 68-72 nucleotides mapping to full-length tRNAs. Interestingly, 18 out of 22 mitochondrial tRNA species were detected in all the age time points, while only 4 cytosolic full-length tRNAs were detectable. All the identified full-length tRNAs were found to be increased in 22-month-old β-cells (**Suppl. Figure 3.e**). This increase may be caused by the upregulation of tRNA transcription and/or to the modulation of specific nucleotide modifications affecting library preparation.

### miRNA and tRF modulation in other aging tissues

We next assessed whether the observed deregulation of miRNAs and tRFs is specific for β-cells or is shared with other metabolic tissues. For this purpose, selected miRNAs and tRFs identified by RNA-sequencing were measured by qPCR in FACS-sorted β-cells, liver, skeletal muscle (SM), adipose tissue (AT) and in medial pre-frontal cortex (mPFC), a brain region involved in age-related cognitive decline (**Figure 3**). None of the tested sncRNAs showed a universal pattern in response to ageing. We found that the inflammatory marker miR-146a-5p was mainly induced in β-cells (confirming our sequencing data), in AT and mPFC (**Figure 3.a**). As previously reported in elderly rat islets [20], miR-181a-5p was decreased in FACS-sorted mouse β-cells but also in SM, and a similar tendency was observed in liver (**Figure 3.b**). The levels of miR-143-3p and miR-145-5p were strongly induced in aging β-cells, an effect that was not observed in any other tissue except for mPFC in which a smaller increase of miR-143-3p was detected (**Figure 3.c-d**). The expression of miR-34 family has been extensively associated to p53 signaling in cancer and ageing tissues [32]. A progressive induction of miR-34a-5p was observed in most analyzed tissues, including SM, AT and mPFC. In FACS-sorted β-cells, miR-34a-5p tended to increase at 12-months and to decrease at 22-months of age (**Figure 3.e**). Concerning the tRFs, we found that the induction of 5’tRF^Gly(GCC)^ was specific to β-cells (**Figure 3.f**). In contrast, modulation of the mitochondrial tRFs 5’tRF^Ser(GCT)^ and itRF^Leu(TAA)^ was also detected in other metabolic tissues (**Figure 3.g-h**).

**FIGURE 3.**
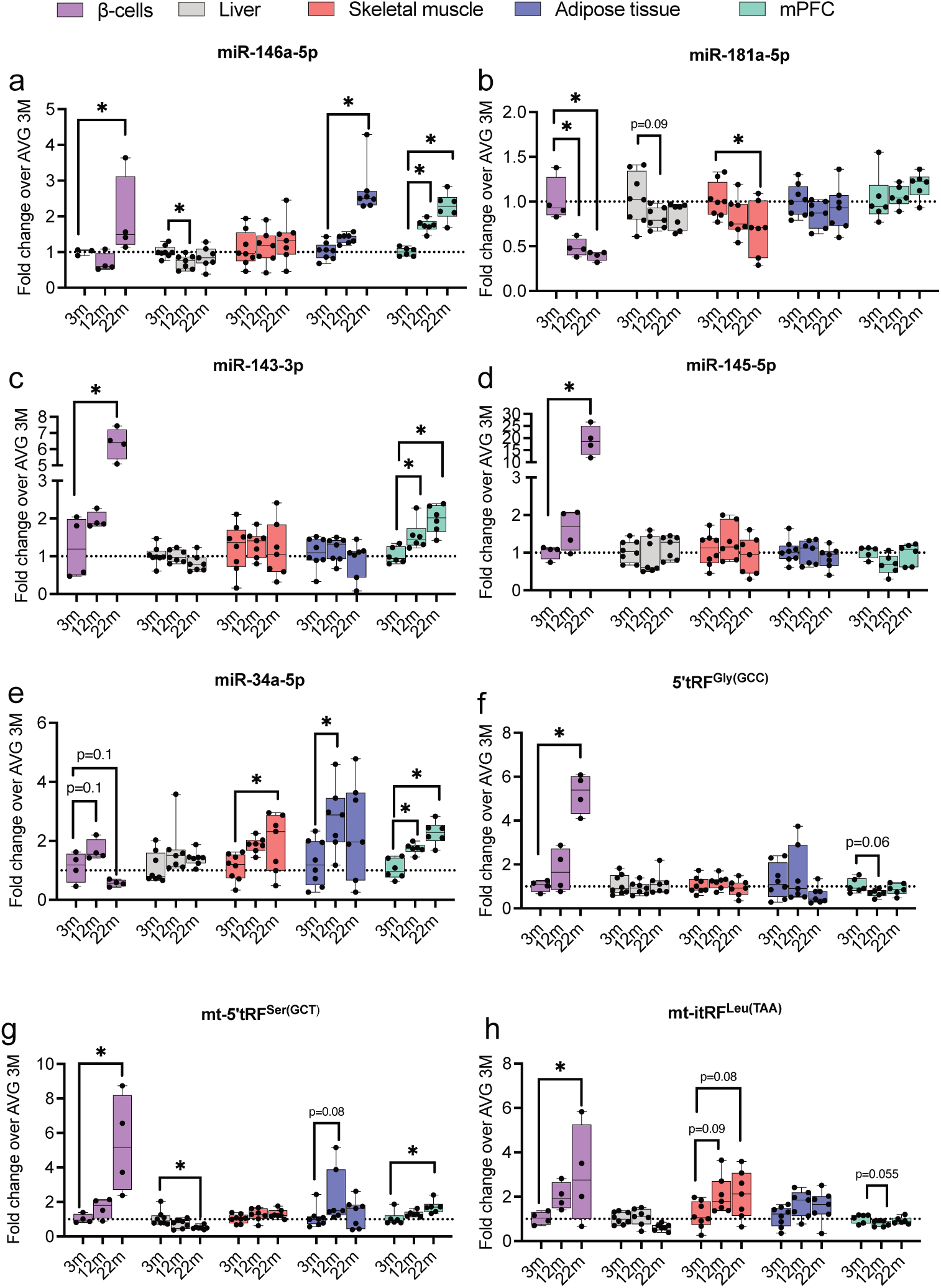
miRNA and tRF levels in aged metabolic tissues. FACS-sorted β-cells (purple), liver (grey), skeletal muscle (red), adipose tissue (blue) and medial pre-frontal cortex (mPFC, green) were isolated from 3-, 12- and 22-month-old C57BL/6J male mice. RNA was extracted and a-e) miRNA and f-h) tRF levels were measured by qPCR, normalized on let-7a-5p and let-7b-5p and expressed as fold change over the average value of the specific tissue from 3-month-old mice. *p<0.05 by one-way ANOVA, Dunnett’s post hoc test.

### miRNA and tRF profiling of senescent β-cells

Since senescence is closely associated with aging, we next investigated if miRNA and tRF profiles of senescent β-cells overlap with those observed in β-cells of aged mice. For this purpose, we performed a small RNA sequencing of senescent β-Gal^+^ β-cells isolated from 7/8-month-old mice [8]. By comparing the expression profiles of β-Gal^+^ and β-Gal^-^ β-cells (**Figure 4.a)**, we identified 41 miRNAs differentially expressed in senescent β-cells, most of them (36) being upregulated. Among the miRNAs affected by senescence, 19 were also modulated during aging (**Figure 4.b**). We found that the induction of miRNAs linked to islet inflammation including miR-146a-5p and miR-142-3p/-5p was comparable and, in some cases, even greater in senescent β-cells than in 22-month-old β-cells. Notably, the level of miR-155-5p, which is also associated to inflammation, was only increased in senescent β-cells (**Figure 4.b and d**). This may indicate that an early induction of inflammatory signals promotes the acquisition of a senescent phenotype in a subset of β-cells. The miR-143/miR-145 cluster, which is associated with insulin resistance [33], was similarly induced in β-Gal^+^ β-cells and in β-cells of 22-month-old mice. In contrast, the levels of miR-181a-5p and miR-181b-5p that decrease with aging are increased in senescent β-cells (**Figure 4.b and d**). In rat β-cells, miR-181b has been shown to target the mRNAs of glycerol-3-phosphate dehydrogenase 2 *(Gpd2)* and malate dehydrogenase 1 *(Mdh1*), two key genes involved in glucose metabolism and insulin secretion and also sirtuin 1 (*Sirt1*) and transforming growth factor β receptor 1 (*Tgfbr1*) [34]. Interestingly, the increased levels of miR-181 observed in senescent β-cells correlates with the reduction of Sirt1 that have been previously associated to the acquisition of a senescent phenotype and premature aging in β-cells and in other cells [5, 35]. Changes in 22 miRNAs were found to be specific to senescent β-cells and not linked to aging (**Figure 4.c**). Among those, the induction of miR-184-3p (**Figure 4.d**), a miRNA previously described to mediate β-cell apoptosis and to reduce proliferation [36, 37]. The level of this miRNA has been reported to be strongly decreased in obesity and T2D, probably as a result of an adaptive process to compensate for insulin resistance. While aging was associated with profound changes in the tRF profile, β-cell senescence had only a modest impact on tRF levels (**Figure 4.e-f**) and mitochondrially-encoded tRFs were not affected in β-Gal^+^ β-cells.

**FIGURE 4.**
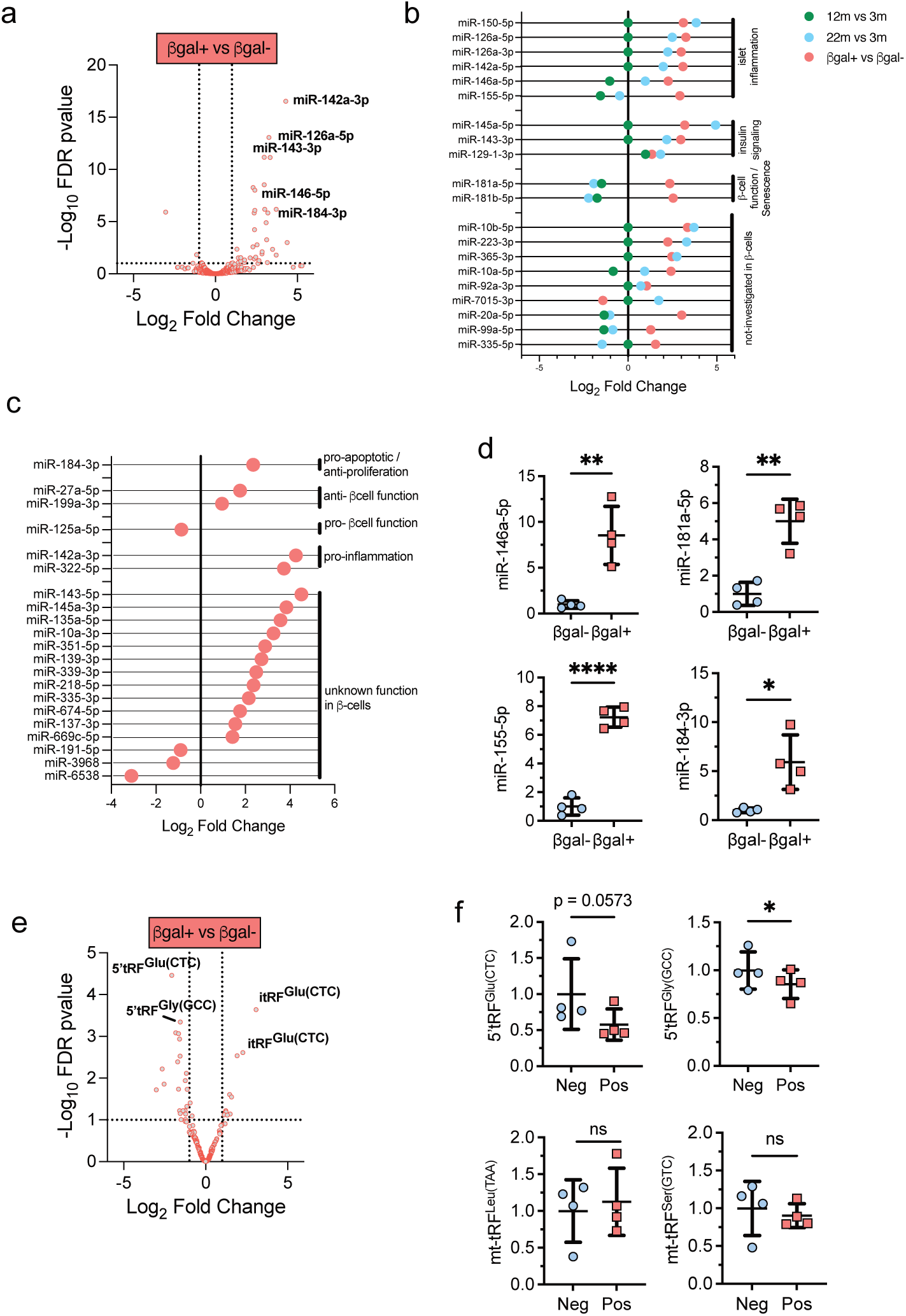
Senescence induced miRNA and tRFs changes in β -cells. Small RNAseq on FACS-sorted SA-βGal positive versus SA-βGal negative β-cells isolated from 7/8-month-old male mice. a) Volcano Plot shows differentially expressed miRNAs in senescent (βGal^+^) versus control (βGal^-^) cells (FDR p-value <0.1, FC>1.5). b) Schematic representation of specific miRNA expression changes in aged or senescent β-cells and grouped by cellular functions, if known in the literature. c) Schematic representation of miRNA changes in senescent β-cells and clusters by cellular functions. d) miRNA changes confirmed by qPCR. miRNA levels were normalized on miR-7a-5p and let-7b-5p and expressed as fold change over the average value of βGal^-^ cells. *p<0.05 by Student’s t test. e) Volcano Plot shows differential tRF levels in senescent (βGal^+^) versus control (βGal^-^) cells (FDR p-value <0.1, FC>1.5). f) tRF changes confirmed by qPCR. tRF levels were normalized on miR-7a-5p and let-7b-5p levels and expressed as fold change over the average value of βGal^-^ cells. *p<0.05 by Student’s t test.

### Inflammatory conditions drive age-associated miRNA and tRF modulation

Since inflammation seems to drive aging and senescent phenotypes, we next investigated if *in vitro* incubation of mouse islet cells with pro-inflammatory cytokines could recapitulate the modulation of sncRNAs observed in β-cells. Since it is challenging to reproduce on a short period of time a process that can take years to develop, we included a wash-out period of 3 days to permit to the aging/senescent-mimicking phenotype to establish. Dispersed mouse islet cells were exposed for 48h to a mix of pro-inflammatory cytokines (30 ng/ml IFNγ, 10 ng/ml TNFα and 1 ng/ml IL-1β) followed, or not, by 3 days wash out period to assess early and late changes (**Figure 5.a**). *p21Cis,* an early senescent marker, and *IL6* expression levels were induced already after 48h exposure to the pro-inflammatory cytokines, while *Bambi* and *Serpine1* were only induced at a later stage (**Figure 5.b-e**). The levels of miR-142-3p, miR-146-5p, miR-181a-5p and 5’tRF^Gly(GCC)^ were already altered upon 48h cytokine treatment (**Figure 5.j-l,o**). In contrast, other miRNAs and tRFs such as miR-10b-5p, miR-143-3p, miR-145-5p, miR-184-3p, mt-5’tRF^Ser(GCT)^ and mt-itRF^Leu(TAA)^ were only significantly modulated at the end of the wash-out period (**Figure 5.f-i and n-o**). Taken together, these findings indicate that prolonged exposure to pro-inflammatory cytokines can, at least partially, reproduce sncRNA changes occurring in senescence and aging β-cells.

**FIGURE 5.**
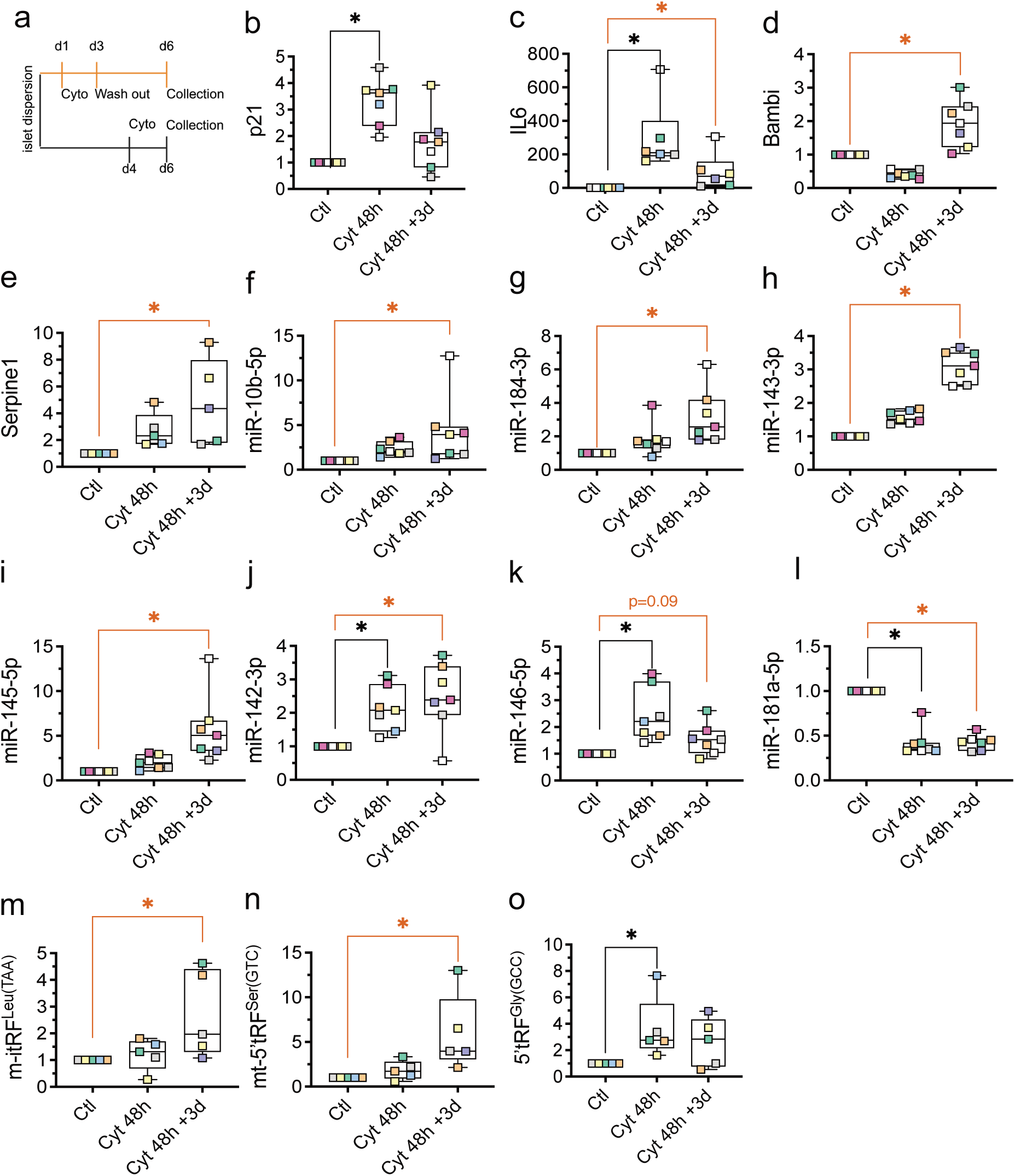
In vitro exposure of islet cells to pro-inflammatory cytokines induced miRNA and tRF changes related to aging and senescence. a) Experimental design. Mouse islet cells were isolated and dispersed. The next day (d1), a group of dispersed cells (Cyt48h+3d) were exposed to pro-inflammatory cytokines (30 ng/ml IFNγ, 10 ng/ml TNFα and 1 ng/ml IL-1β) for 48h and media was changed on d3 to normal media for 72h (washout period). In parallel, another group of cells were first incubated in normal media for 4 days and exposed to pro-inflammatory cytokines for the last 48h (Cyt 48h). Control (Ctl) cells were cultured with normal media for 6 days. On d6, RNA was extracted at the same time for all cell conditions. B-d) mRNA, e-k) miRNA and l-n) tRF levels were measured by qPCR, normalized on HPRT (mRNA) or let-7a-5p and let-7b-5p levels (miRNAs and tRFs) and expressed as fold change over the control group. *p<0.05 by one-way ANOVA, Dunnett’s post hoc test.

### Effect of aging on the small RNA profile of islet-resident macrophages

Islet resident macrophages display a peculiar inflammatory-like phenotype and interact closely with β-cells, sensing and influencing their functional and metabolic state. These plastic cells respond to changes in islet micro-environment by transcriptional and metabolic remodeling to support tissue adaptation. To assess whether the activation state of iMACs is modified during aging, we optimized the FACS-sorting protocol to collect in parallel iMACs and β-cells from our mouse aging model (**Suppl. Figure 1.c-d**). As reported for other macrophage populations [11], we found that the proportion of iMACs per islet cells increases during aging (**Figure 6.a**). We then investigated age-dependent miRNA changes in iMACs potentially influencing their activation state. We found 8 miRNAs in 12-month and 16 miRNAs in 22-month-old mice differentially expressed compared to young mice (FDR pvalue <0.1, FC>1.5; **Figure 6.b-c**). Functional enrichment analysis showed an overrepresentation of miRNAs involved in cell division, migration and immune cell differentiation (**Figure 6.d**). Overall, we observed an increase in the level of anti-inflammatory miRNAs and a decrease of those with pro-inflammatory actions (**Figure 6.e**). The most pronounced age-dependent effect was observed for miR-206-3p that was induced >300 folds at 22 months. This miRNA has been shown to inhibit inflammation, targeting the TLR4/NF-κB/NLRP3 pathway [38]. Upregulation of miR-126a-5p and miR-126a-3p was already detected at the age of 12 months and was maintained at 22 months. MiR-126a has been reported to induce the switch from pro-inflammatory to anti-inflammatory macrophage phenotype, and to promote efferocytosis, a phagocytic mechanism by which tissue macrophages eliminate apoptotic cells [39]. The pro-inflammatory miR-155-5p was downregulated at 22 months. The modulation of these miRNAs was confirmed by qPCR (**Figure 6.f-g**).

**FIGURE 6.**
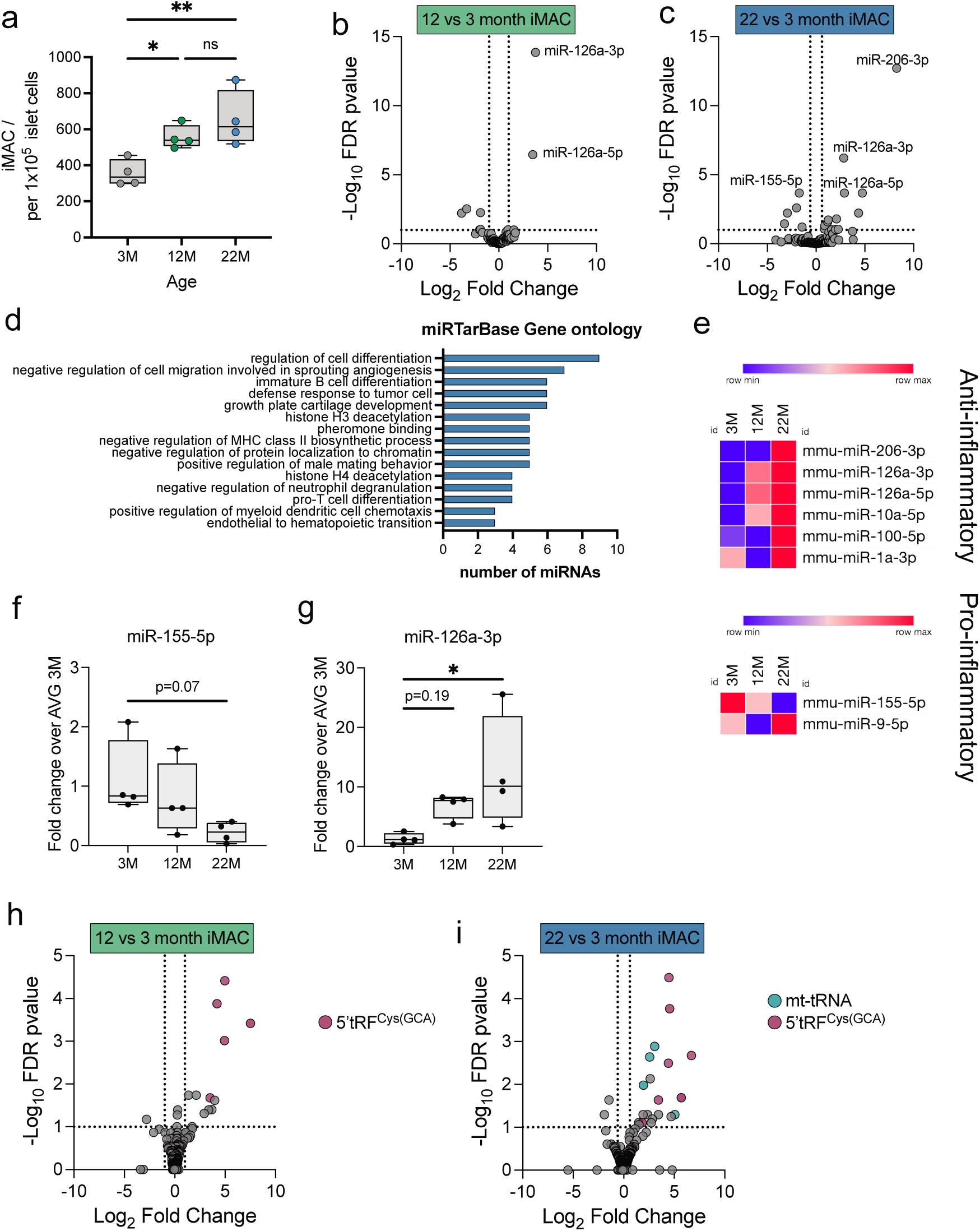
Effect of aging on islet-resident macrophage miRNA and tRF profiles. Islet-resident macrophages (iMACs) were FACS-sorted from young adult (3-month), middle-aged (12-month) and elderly (22-month) C57BL/6J male mice. a) Number of FACS-sorted iMACs increased with age. b-c) Volcano Plots show differentially expressed miRNAs in 12-month (b) or 22-month (c) versus 3-month-old iMACs (FDR p-value <0.1, FC>1.5). d) miRTar Base Gene Ontology enrichments associated with miRNA downregulation of 22-month-old iMACs. e) Heatmap plots showing miRNAs significantly deregulated in 22-month-old iMACs that were previously reported in the literature to be associated with macrophage anti-inflammatory or pro-inflammatory states. f-g) miRNA changes confirmed by qPCR. miRNA levels were normalized on miR-16-5p and expressed as fold change over the average value of 3-month-old iMACs. *p<0.05 by one-way ANOVA, Dunnett’s post hoc test. h-i) Volcano Plots show differentially expressed tRFs in 12-month (h) or 22-month (i) versus 3-month-old iMACs (FDR p-value <0.1, FC>1.5). Purple circles correspond to tRFs derived from nuclear encoded tRNAs, while turquoise circles refer to tRFs derived from mitochondrial encoded tRNAs.

Changes in the iMAC tRF profile occurring during aging revealed the induction of several 5’tRFs derived from the cytosolic tRNA^Cys(GCA)^ and from different mitochondrial tRNAs (**Figure 6.h-i**). The modulation of 5’tRF^Glu(CTC)^ in iMAC has been recently shown to be linked to their activation state [9], however the function of other tRFs has not been investigated so far in iMAC.

## DISCUSSION

This study aimed to perform a comprehensive profiling of sncRNA changes occurring in aging and senescent FACS-sorted β-cells and islet-resident macrophages (iMACs). These datasets complement and extend previous research into changes in β-cell functions and in gene expression associated with aging and cellular senescence.

We confirmed several miRNA changes previously observed in aged rat islets [20] within sorted mouse β-cells. Specifically, miR-181a-5p was decreased in 22-month-old mice, while miR-124-3p, miR-129-1-3p, miR-383-3p and miR-488-3p were upregulated. Additionally, we observed a trend for a decrease in miR-34a-5p levels in elderly β-cells, although this did not reach statistical significance (p=0.1). Xu and colleagues also examined miRNA deregulation in brain, kidney and liver of 22-month-old mice, observing changes in miR-124-3p and miR-29 family (which was also similarly deregulated in our aged β-cell dataset). Their investigation in mouse embryonic fibroblasts (MEF) revealed that p53 drives the expression of miR-124-3p and miR-29a/b/c-3p, which target p21 antagonist *Ccna2*, thereby promoting cellular senescence [40]. This suggests that miR-124-3p and miR-29 family members may play a role in the development of cellular senescence during aging.

Furthermore, we identified several miRNAs deregulated in aging and/or senescent β-cells that are known to be modified under diabetic conditions. For example, miRNAs linked to inflammation that are increased in islets of T1D models, such as miR-142-3p, miR-142-5p, miR-146a-5p, miR-150-5p and miR-155-5p were also elevated in aging and/or senescent β-cells. Both aging and senescence are associated to chronic low-grade inflammation called « inflammaging » and « senoinflammation ». Some of these changes could be recapitulated by exposing *in vitro* mouse islet cells to pro-inflammatory cytokines. Among them, miR-146a-5p which is known to regulate inflammation not only in β-cells [28, 29] but also in other tissues and has been named inflammaMir [41].

Other miRNAs, including the miR-132-3p/miR-212 cluster, the miR-29 family, miR-204-5p and miR-802-3p, have previously been reported to be deregulated in islets of various obese and T2D models, where they impact insulin secretion [36, 42–46]. In elderly humans, β-cells exhibit impaired insulin secretion in response to glucose. In rodents, conflicting findings have been reported regarding glucose-stimulated insulin secretion in aged mouse islets, although increased basal insulin secretion is generally observed (reviewed in [2]). Elevated basal insulin secretion is increasingly seen as an early sign of β-cell dysfunction that predisposes to insulin resistance [47, 48].

As evidenced in the heatmap in Figure 1, most miRNAs promoting proliferation, including miR-181a-5p, miR-181b-5p and miR-338-3p [20, 34] were reduced, while miR-132-3p was upregulated. Additionally, miRNAs that inhibit β-cell maturation (miR-15b-5p, miR-181a/b) were downregulated, while those promoting maturation (miR-129, miR-132-3p, miR-204-5p, miR-344-3p, miR-383-5p, miR-415a) were upregulated (Figure 1). Together, these findings suggest that 22-month-old β-cells exhibit a fully mature phenotype with limited proliferative capacity. The situation in senescent β-cells is less clear. While miR-184-3p, a known inhibitor of β-cell mass expansion [37] was increased, miR-181a-5p and miR-181b-5p, which promote proliferation, were also elevated. This suggests that the reduced proliferative ability in aged and senescent β-cells may arise from partially distinct mechanisms.

Thanks to the advancement of sequencing technologies, variations in miRNA sequence were identified to be highly represented. IsomiRs may include nucleotide additions, deletions and substitutions mediated by different mechanisms. While nucleotide additions in the 5’ region and variations in miRNA seed sequence may directly affect target recognition, 3’ insertions have been reported to have an impact on the stability of several miRNAs [15, 19]. The modulation of the level of specific isomiR can regulate β-cell functions [16]. However, the biogenesis and roles of isomiRs are only beginning to be unveiled and the functional implications of the changes in isomiR levels observed in aged β-cells remain to be established.

Fragments resulting from tRNA cleavage represent a class of functional sncRNAs involved in islet cell homeostasis [49] but their role in cellular senescence and aging remains poorly understood [50]. The biogenesis of cytosolic 5’tRF has been identified as a stress-response mechanism in T2D and a pathogenic feature underlying TRMT10A monogenic diabetes [18]. We observed an upregulation of the 5’tRF, derived from Gly-tRNA^(GCC)^ in aged β-cells, which is also increased in the islets of NOD mice during early T1D development [17]. Overexpression of this tRF increases β-cell apoptosis, partly by reducing the level of the anti-apoptotic protein Bcl-XL [17].

In our dataset, we noted a global upregulation of mitochondrial tRFs (mt-tRFs) in 22-month-old β-cells. Notably, several mt-tRF have been previously found to increase in the islets of T1D models [17], but to decrease in the islets of T2D models [31]. Among the upregulated mt-tRFs in aged β-cells were mt-5’tRF^Ser(GCT)^ and mt-itRF^Leu(TAA)^. The former, was previously found to be upregulated in the islets of prediabetic NOD mice and in response to pro-inflammatory cytokines [17], although its function in β-cells is yet to be determined. The internal tRF mt-itRF^Leu(TAA)^ is decreased in the islets of T2D models, resulting in a reduction in mitochondrial respiration and glucose-induced insulin secretion. In contrast, increased levels of mt-itRF^Leu(TAA)^ favored glucose-induced insulin secretion [31]. Interestingly, we observed a trend toward decreased mt-itRF^Leu(TAA)^ levels in skeletal muscle of aged mice. Previous work indicated that the downregulation of mt-itRF^Leu(TAA)^ in skeletal muscle increases insulin sensitivity [51]. Thus, the reduction of mt-itRF^Leu(TAA)^ in aged mice may contribute to insulin resistance.

The increase in tRF levels may result either from enhanced tRNA synthesis or greater cleavage activity [52]. Although full-length tRNAs are heavily modified and typically challenging to detect using standard RNA sequencing protocols, our sequencing settings allowed to detect molecules up to 80 nucleotides, including some full-length tRNAs. Most of the detected tRNAs were mitochondrially encoded, possibly because they are less modified than cytosol tRNAs [53]. Importantly, all detected mitochondrial tRNAs were found to be upregulated in 22-month-old β-cells. Our experimental design cannot determine whether this elevation is due to increased tRNA synthesis or changes in post-transcriptional modifications that make full-length tRNAs more detectable.

An additional indication of mitochondrial ncRNAs deregulation in aging comes from our unbiased dataset annotation. We found an increase in fragments derived from mitochondrial ribosomal RNAs (rRNAs) and mitochondrial-coding genes in β-cells from aged mice. Interestingly, rRNA-derived fragments have been recently shown to possess regulatory properties [54]. Future studies investigating these poorly characterized sncRNAs may uncover new aspects of the molecular mechanisms underlying dysfunction in aging and senescent β-cells. This emphasizes the importance of a comprehensive, sequence-centric analysis with appropriate filtering strategies to gain a complete picture of the cellular sncRNome.

Finally, our study provides the first characterization of the iMAC sncRNA profile during aging. While the critical role of iMACs in establishing an appropriate functional β-cell mass [55] and in regulating the plasticity of islet stress responses during obesity and T2D has been documented [9, 10, 56], their involvement in maintaining islet homeostasis during aging remains poorly understood. Our data show that iMACs from aged mice exhibit reduced expression of pro-inflammatory miRNAs and a concomitant increase in miRNAs with anti-inflammatory and pro-phagocytosis functions. Although these findings will need to be validated through transcriptional and functional analysis, they likely reflect a phenotypic shift in iMACs aimed at preserving islet homeostasis in aging animals.

## CONCLUSIONS

This study aimed at providing a comprehensive overview of sncRNA changes during aging in pancreatic islet cells. We provide detailed analysis of miRNAs, isomiRs, piRNAs and tRFs, along with differential expressions of all small non-coding sequences unambiguously mapping to one genomic location in 12-and 22-month-old murine β-cells. We observed changes related to β-cell inflammation and mitochondrial functions. Functional studies remain to be performed to better understand the role of sncRNA deregulation in the complexity of aging process in both mice and humans. However, these datasets are complementary with transcriptomic and proteomic data previously published [1, 5, 57] and all together, may help predicting individuals who will age healthy versus the one predisposed to develop metabolic diseases like diabetes.

## ACKNOWLEDGEMENTS

The authors thank Rémy Klein for technical help, Danny Labes and Mariela Castleblanco from the Flow Cytometry Plateform (University of Lausanne) for β-cell and iMAC cell sorting, and Nicolas Guex and Christian Iseli from the Bioinformatics Competence Centre (University of Lausanne) for RNAseq analyses. This work was funded by Swiss National Science Foundation (grant 310030-219252) to RR and by the Transition Grant from the faculty of Biology and Medicine, University of Lausanne to CC.

## AUTHOR CONTRIBUTION

RR, CC and CG designed the study, conducted data analysis, and wrote the manuscript. CC, CG, JP, VM performed experiments and analyzed the data. EM conducted sncRNA-seq data bioinformatical analyses. SBW and CA provided senescence β-cells and scientific expertise. EA, DF, FD and GS provided isomiRs bioinformatical analyses and scientific expertise. RR and CC provided financial support. All authors contributed to the manuscript preparation and approved the final version.

## ETHICS APPROVAL

This study was approved by the Swiss Research Councils and Veterinary Offices (authorization number VD2495x4) and by Animal Care and Use Committee from the Joslin Diabetes Center (Protocol number 2022-01).

## ABBREVIATIONS

AT: Adipose tissue
FACS: Fluorescence-Activated Cell Sorting
FC: fold change
iMACs: Islet-resident macrophages
IFNγ: interferon γ
IL-1β: interleukin 1β
isomiRs: isoforms of miRNAs
mPFC: medial pre-frontal cortex
miRNAs: microRNAs
mTOR: mammalian target of rapamycin
NOD: non-obese diabetic mice
piRNAs: PIWI-interacting RNAs
RNAseq: RNA sequencing
SA-β-Gal: senescence-associated acidic β-galactosidase
SM: skeletal muscle
sncRNA: small non-coding RNAs
T1D: type 1 diabetes
T2D: type 2 diabetes
TNFα: Tumor Necrosis Factor α
tRFs: tRNA-derived fragments

## SUPPLEMENTARY FIGURES AND FIGURE LEGENDS

**Suppl. FIGURE 1.**
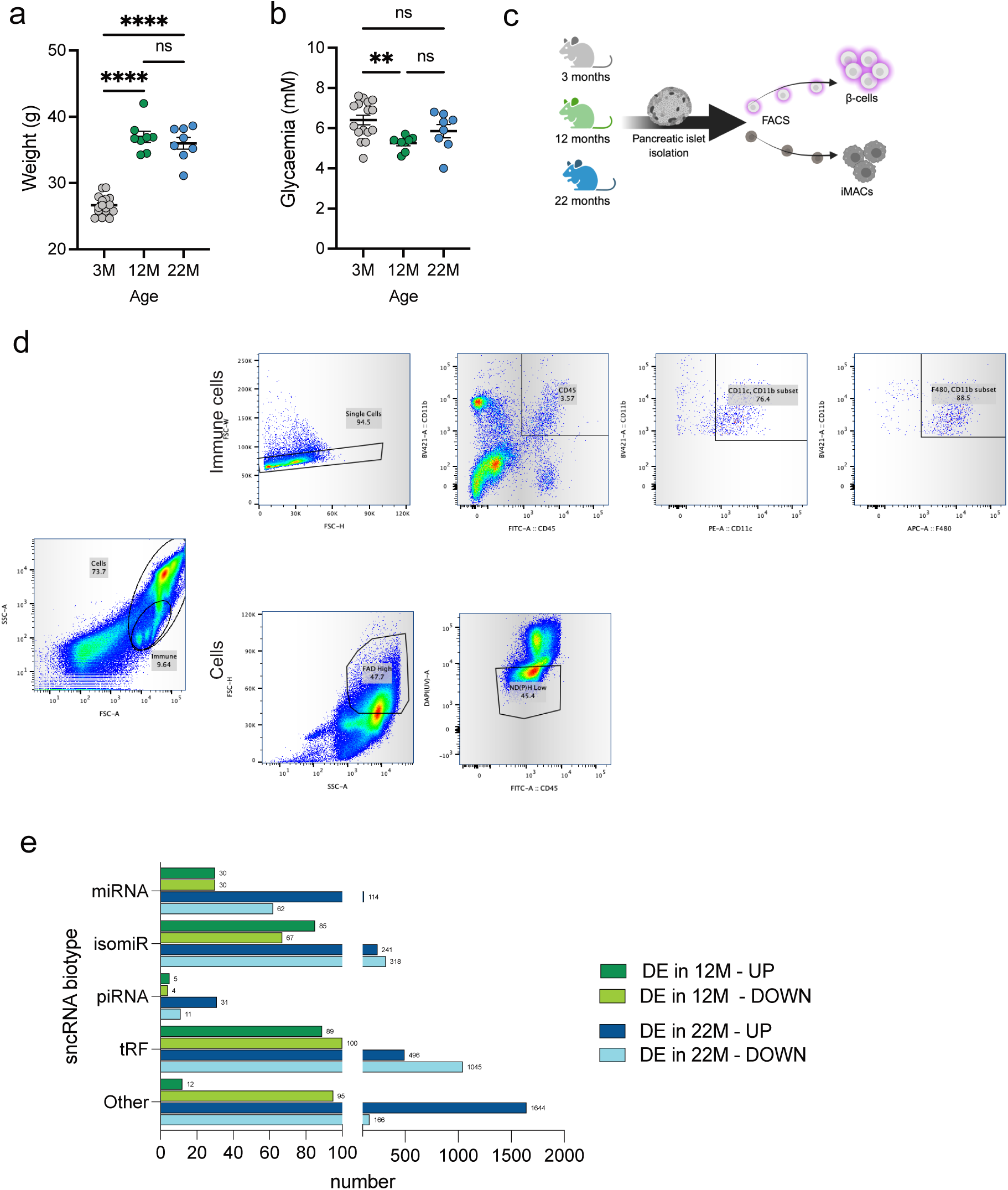
sncRNA profiling in aged β-cells and iMACs. A) Body weight and B) Glycemia of 3-, 12- and 22-month-old mice in fed state. C) Experimental design: Pancreatic islets were isolated from 3-, 12- and 22-month-old mice, dispersed and FACS sorted to collect enriched β-cell (99%) and islet-resident macrophage (iMACs) fractions. D) Gatting strategy: Immune cells were sorted based on the expression of CD45, Cd11b, F4/80 and Cd11c markers (top panel) while β-cells were sorted based on autofluorescence (lower panel). E) Differentially expressed (DE) sncRNA biotypes (miRNAs, siomiRs, piRNAs, tRFs and others) in middle-aged (12 months – green bars) and elderly (22 months-blue bars) β-cells compared to young adult (3 months). Dark green/blue bars correspond to increased levels while light green/blue bars correspond to decreased levels of sncRNAs. Fold change ≥1.5 and FDR≤0.1 were considered significant (number of significant changes for each biotype and aged condition is indicated next to the bar).

**Suppl. FIGURE 2.**
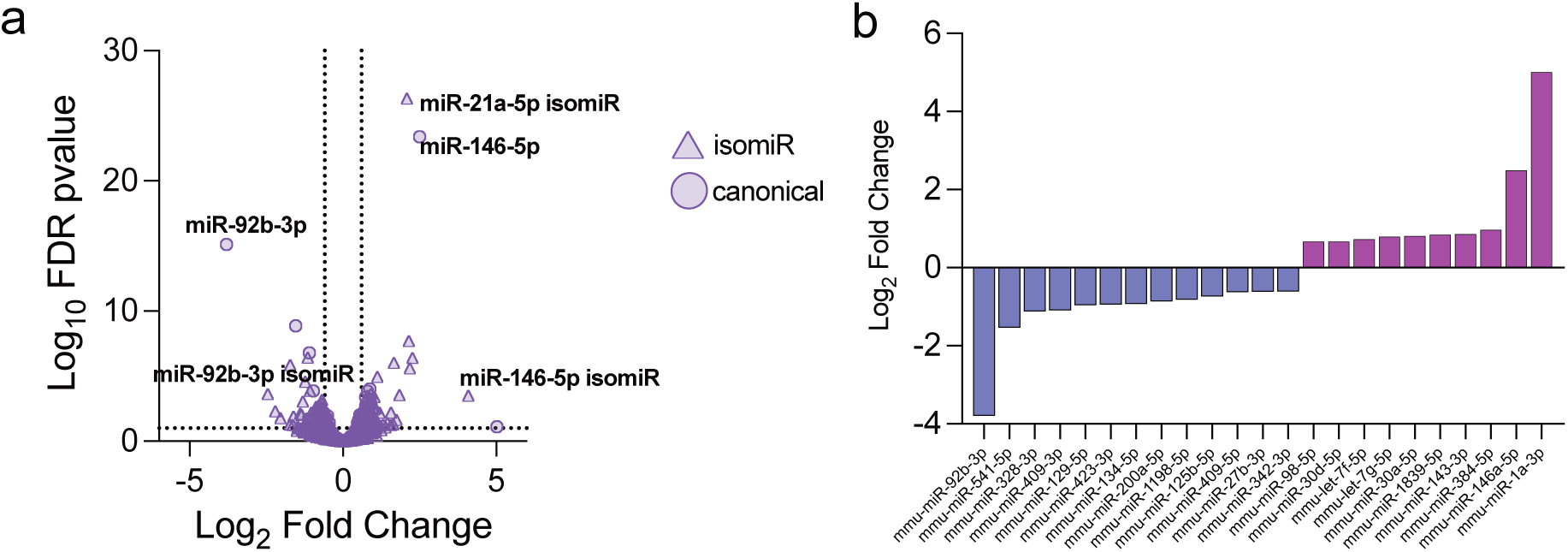
Global increased of mitochondrial RNA transcripts in aged β-cells. Volcano Plot shows differentially expressed canonical miRNA (circles) and isomiRs (triangles) in mouse islets treated for 24h with pro-inflammatory cytokines compared to not treated control (FDR p-value <0.1, FC>1.5).

**Suppl. FIGURE 3.**
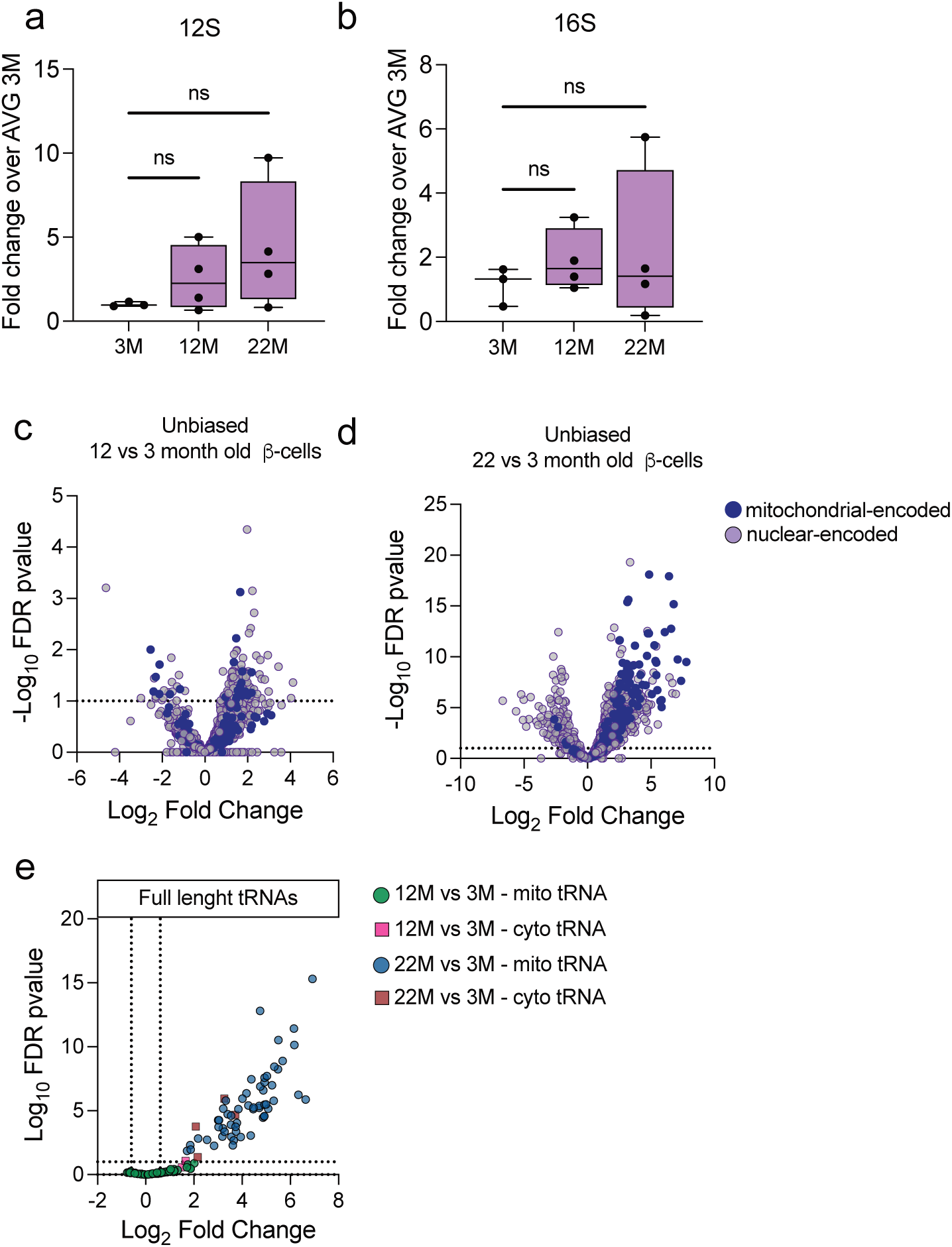
Global increased of mitochondrial RNA transcripts in aged β-cells. A) 12S and B) 16S expressions in FACS-sorted β-cells of 3-, 12- and 22-month-old mice. Ribosomal RNA levels were measured by qPCR, normalized to HPRT and expressed as fold change over the average value of 3-month-old β-cells. C-D) Unbiased analysis of sequences mapping to mitochondrial-(dark purple) or nuclear-encoded (light purple) RNA transcripts in aged C) 12-month-old β-cells or D) 22-month-old β-cells versus young 3-month-old β-cells. E) Differential expressions of full length tRNA transcripts detected in small RNAseq. Circles: mitochondrial-encoded tRNA changes in aged β-cells of 12- (green) or 22- (blue) month-old mice versus young 3-month-old β-cells. Squares: Nuclear-encoded tRNA changes in aged β-cells of 12-(pink) or 22- (red) month-old mice versus young 3-month-old β-cells.

**Suppl. TABLE 1.**
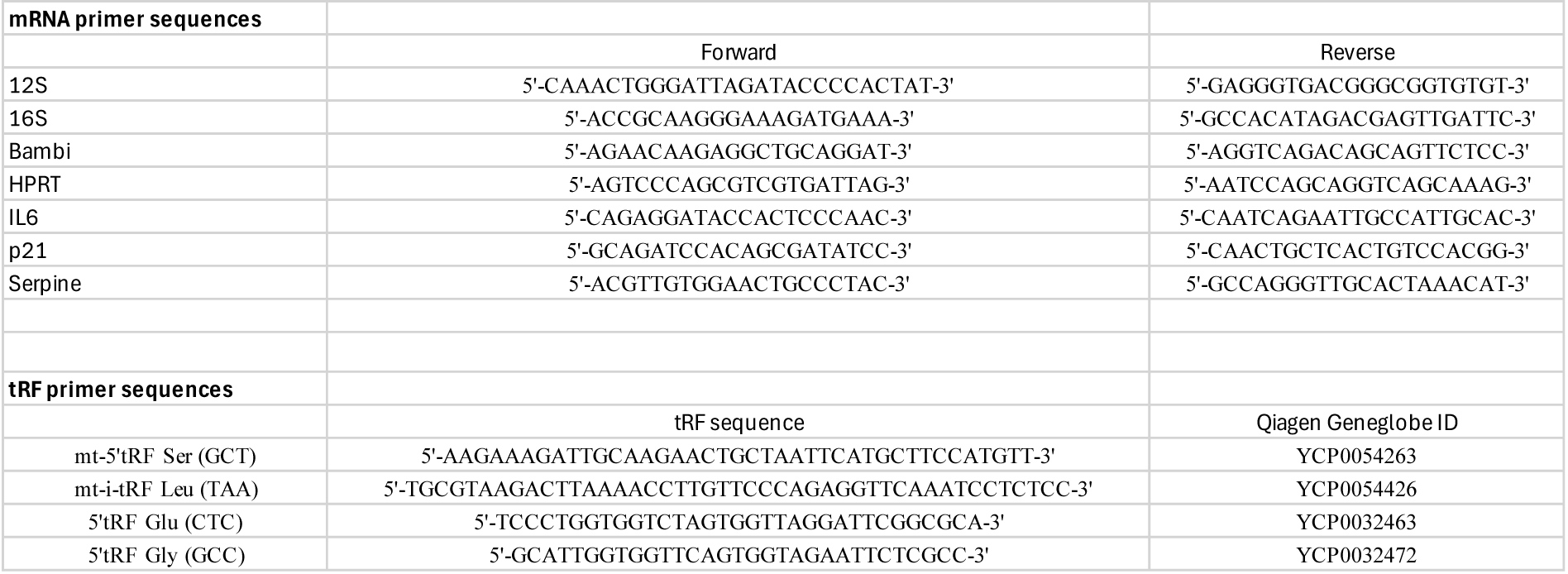
mRNA and tRF primer sequences. Primer sequences and target nucleotide sequences

